# Endocytosis at Extremes: Formation and Internalization of Giant Clathrin-coated Pits Under Elevated Membrane Tension

**DOI:** 10.1101/2022.06.07.495153

**Authors:** Ahmet Ata Akatay, Tianyao Wu, Umidahan Djakbarova, Cristopher Thompson, Emanuele Cocucci, Roya Zandi, Joseph Rudnick, Comert Kural

## Abstract

Internalization of clathrin-coated vesicles from the plasma membrane constitutes the major endocytic route for receptors and their ligands. Dynamic and structural properties of endocytic clathrin coats are regulated by the mechanical properties of the plasma membrane. Here, we used conventional fluorescence imaging and multiple modes of structured illumination microscopy (SIM) to image formation of endocytic clathrin coats within live cells and tissues of developing fruit fly embryos. High resolution in both spatial and temporal domains allowed us to detect and characterize distinct classes of clathrin-coated structures. For the first time, we show that membrane tension induces formation of giant coated pits (GCPs) that can be up to two orders of magnitude larger than the canonical clathrin-coated pits. GCPs take longer to form but their mechanism of curvature generation is the same as the canonical pits. We also demonstrate that GCPs can split into smaller fragments during internalization. Considering the supporting roles played by actin filament dynamics in clathrin-mediated endocytosis under mechanically stringent conditions, we suggest that local changes in the coat curvature driven by actin machinery can drive splitting and internalization of GCPs.

## INTRODUCTION

Clathrin-coated vesicles are the fundamental functioning units of lipid and protein trafficking from the plasma membrane to endosomes (Conner and Schmid, 2003). They are formed by gradual recruitment of heterohexameric clathrin triskelions to the plasma membrane by a diverse set of adaptor proteins that bind to clathrin, membrane, and other adaptor and accessory proteins (Taylor, Perrais and Merrifield, 2011; Kural and Kirchhausen, 2012). Clathrin triskelions assemble into polyhedral cages (100-200nm in diameter) that generate endocytic pockets on the plasma membrane, altogether referred as clathrin-coated pits (CCPs). As the membrane neck linking the CCP to the plasma membrane gets narrower, dynamin is recruited to this region in a burst and drives membrane scission that leads to internalization of a clathrin-coated vesicle (Cocucci, Gaudin and Kirchhausen, 2014).

Clathrin-coated vesicles can originate through different mechanisms. The best characterized mechanism is based on formation of CCPs *de novo*, where the initiation, maturation, and internalization of the CCP is independent and spatially isolated from other clathrin-coated structures (Willy, Ferguson, *et al*., 2021). In the second mechanism, large clathrin lattices found at membrane-substrate adhesion sites, i.e., clathrin plaques, serve as CCP initiation sites (Lampe, Vassilopoulos and Merrifield, 2016; Leyton-Puig *et al*., 2017). It is proposed that highly curved CCPs forming at the edges of relatively flat plaques increase the strain on the clathrin lattice, eventually giving rise to ruptures and breaks that allow growth and internalization of CCPs independently (den Otter and Briels, 2011; Willy, Ferguson, *et al*., 2021).

Membrane tension is a fast, effective and reversible regulator of endocytic clathrin-coated vesicle formation (Ferguson *et al*., 2016, 2017; Willy *et al*., 2017; Djakbarova *et al*., 2021; Willy, Colombo, *et al*., 2021). Increased membrane tension reduces the initiation density (Ferguson *et al*., 2017), and slows down the growth and dissolution rates of endocytic clathrin-coated structures (Ferguson *et al*., 2016). Moreover, internalization of CCPs from the plasma membrane becomes dependent on the forces provided by actin polymerization under increased tension (Boulant *et al*., 2011).

Here, we studied the dynamics of endocytic clathrin-coated structures within live *Drosophila* embryos and cultured cells using super-resolved fluorescence imaging. In the epidermal tissue of late-stage *Drosophila* embryos, we found that clathrin coats can form dome structures that can be to orders of magnitude larger than the canonical CCPs. In cultured cells, these giant coated vesicles (GCPs) were only observed when the membrane tension was increased by hypotonic swelling or cholesterol depletion. Interestingly, GCPs were internalized by splitting into multiple fragments comparable in size to the canonical CCPs, suggesting a new mechanism of endocytic clathrin-coated vesicle formation. Similar to the “rupture and growth” model suggested for CCP formation at the edges of clathrin plaques, we propose a model where GCPs go through splitting events initiated by the forces provided by the actin machinery giving rise to increased strain and breaks on the clathrin coat.

## RESULTS

### Formation and internalization of canonical clathrin-coated pits (CCPs)

Formation, internalization and dissolution dynamics of endocytic clathrin coats have been characterized in living cells by studies utilizing fluorescence microscopy (Cocucci *et al*., 2012; Kural and Kirchhausen, 2012; Cocucci, Gaudin and Kirchhausen, 2014; Aguet *et al*., 2016; Willy, Colombo, *et al*., 2021). In these assays, CCPs appear as featureless diffraction-limited spots as their typical size is smaller than the resolution limit of the conventional fluorescence microscope. Nevertheless, super-resolved fluorescence microscopy allows to elucidate the structural properties of endocytic CCPs, along with their dynamics (Jones *et al*., 2011; Fiolka *et al*., 2012; Li *et al*., 2015). In particular, structured illumination microscopy at the total internal reflection fluorescence mode (TIRF-SIM) has been instrumental in monitoring curvature generation by endocytic clathrin pits within live cells and tissues (Willy, Ferguson, *et al*., 2021). In these acquisitions, the total internal reflection of the excitation beam creates an evanescent field that illuminates the lateral regions of CCPs at a higher intensity compared to the apex, which is further away from the glass substrate. As a result, formation of CCPs is marked by a characteristic “ring” pattern (100-200nm in diameter) under super-resolution imaging (Willy, Ferguson, *et al*., 2021) (Figure 1A, B). Reducing the incidence angle of the excitation beam enables increasing the penetration depth of the illumination field, i.e., approaching the structured illumination microscopy at the grazing incidence mode (GI-SIM), and imaging deeper inside the cells with enhanced spatiotemporal resolution (Guo *et al*., 2018). Here, we monitored formation of *de novo* CCPs in SUM-159 cells genome edited to express AP2-EGFP (Aguet *et al*., 2016) using high numerical aperture (NA) TIRF-SIM and GI-SIM imaging simultaneously to follow the growth of the coat in the axial dimension (i.e., along *z*) while monitoring curvature generation in real time. We found that the ring pattern arose in the TIRF-SIM channel soon after the nucleation of the coat (Figure 1B - blue boxes). However, due to the longer penetration depth of GI-SIM, the same pattern was not visible in this channel in this time window. As the coat grew further and matured into a pit, the ring pattern was observed in both TIRF-SIM and GI-SIM channels (Figure 1B - green boxes). Altogether, these results demonstrate that the coat is highly curved even at the early stages of clathrin pit formation as proposed earlier (Chen *et al*., 2019; Willy, Ferguson, *et al*., 2021). Interestingly, at later stages, the ring pattern disappeared in the TIRF-SIM channel but persisted in the GI-SIM channel, demonstrating the inward movement (along the axial dimension) and internalization of the clathrin pit prior to its uncoating (Figure 1B - red boxes, Figure 1C). It was previously shown that the inward movement of endocytic CCPs during their internalization is dependent on actin polymerization (Ferguson *et al*., 2017). Moreover, it has been shown that actin-driven displacement of clathrin coats (away from the cell surface) gets longer with increasing membrane tension (Ferguson *et al*., 2017), in accord with models suggesting that actin polymerization pushing the clathrin coats increases with the increasing mechanical load (Kaplan *et al*., 2022).

**Figure 1.**
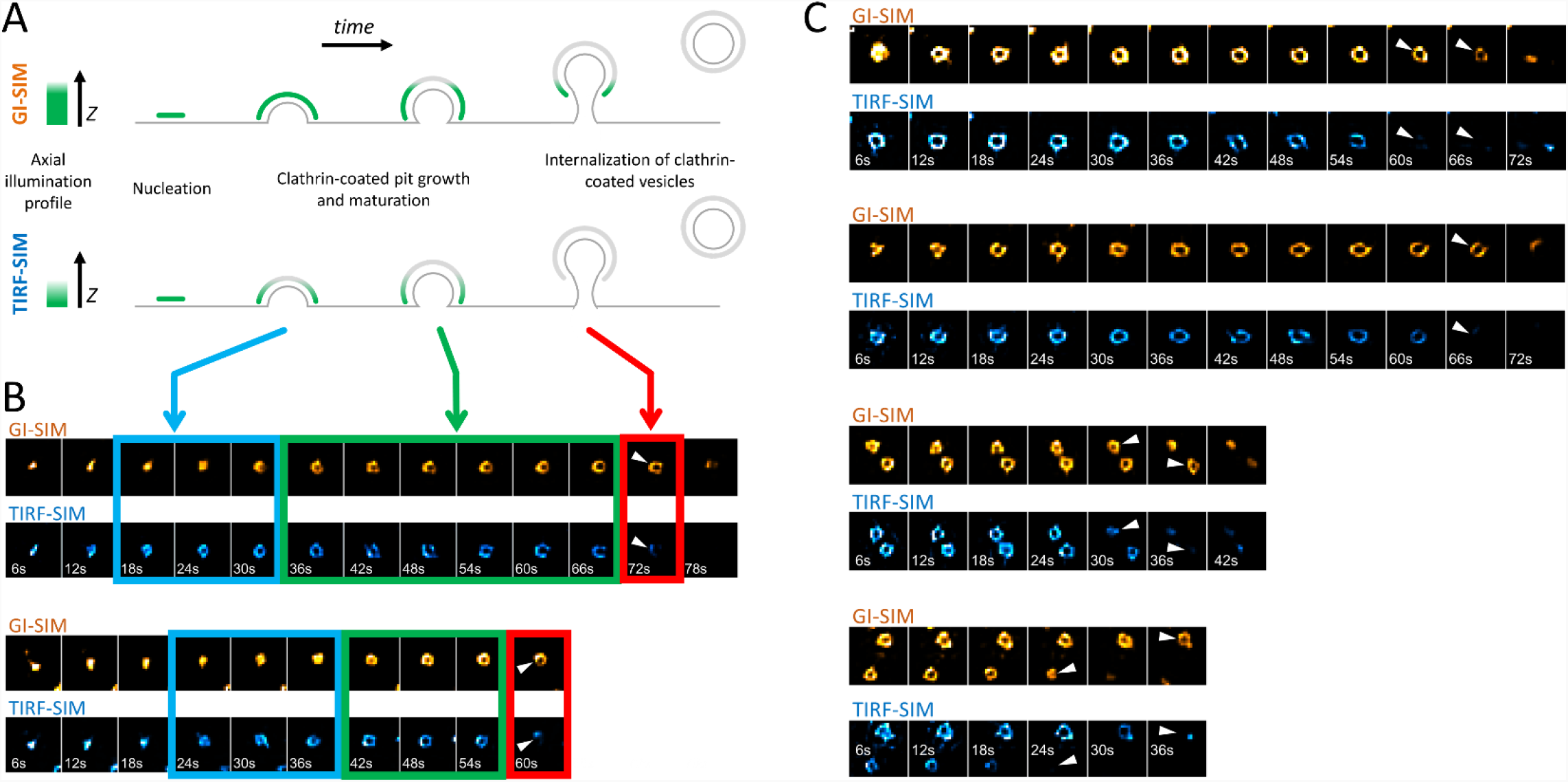
Formation and internalization of “canonical” clathrin-coated pits (CCPs) as detected by simultaneous high NA TIRF-SIM & GI-SIM imaging in living cells. **(A)** GI-SIM offers longer penetration depth, i.e., fluorescence excitation along the *z*-dimension, compared to TIRF-SIM. As invagination of the endocytic pit increases, the apex of the clathrin coat exits the excitation field of TIRF-SIM earlier, resulting in emergence of the ring pattern (blue boxes in B). As the coat matures into a pit, it is observable as a ring in both channels (green boxes in B). While the coat is internalized as a whole, it leaves the TIRF field but still can be observed in the GI-SIM channel (red boxes in B). **(B)** Montages show formation and internalization of two CCPs as imaged by GI-SIM and TIRF-SIM simultaneously in SUM159 cells genome edited to express AP2-EGFP. **(C)** More examples displaying the inward movement of CCPs during internalization. Arrowheads mark disappearance of the clathrin pits from the TIRF-SIM channel while they are still observable by GI-SIM. Each region of interest in B and C, 1*µ*m x 1*µ*m.

### Internalization of clathrin-coated vesicles under high mechanical load

Actin dynamics provide the energy required for internalization of clathrin-coated vesicles under mechanically stringent conditions that slow down completion of clathrin coats (Boulant *et al*., 2012; Ferguson *et al*., 2017; Kaplan *et al*., 2022). For instance, uptake of viral particles larger than the canonical clathrin pits take longer than the time required for internalization of smaller cargo molecules, and necessitate a “push” provided by actin polymerization that moves the clathrin coat away from the plasma membrane (Cureton *et al*., 2010). In certain cell types, the adhesion between the plasma membrane and the substrate slow down the endocytic machinery and result in the formation of clathrin plaques, large and flat clathrin lattices that are longer lived than the canonical CCPs (Batchelder and Yarar, 2010; Ferguson *et al*., 2016; Lampe, Vassilopoulos and Merrifield, 2016). In three-dimensional fluorescence live cell imaging assays, clathrin plaques appear as bright and long-lived puncta localized at cell-substrate contact sites (Figure 2A, B). The edge regions of clathrin plaques are known as actin-dependent endocytic hubs (Lampe, Vassilopoulos and Merrifield, 2016; Leyton-Puig *et al*., 2017), where local mismatches in coat curvature are proposed to give rise to ruptures in the clathrin lattice and allow individual CCPs to split from the plaque and internalize independently (den Otter and Briels, 2011; Willy, Ferguson, *et al*., 2021) (Figure 2C). It was proposed that polymerization of actin filaments generates the “pushing” force required for internalization of plaques from the plasma membrane (Saffarian, Cocucci and Kirchhausen, 2009). Indeed, we found that the growth and dissociation dynamics of clathrin plaques (Ferguson *et al*., 2016, 2017; Willy *et al*., 2017) reduce in SUM-159 cells when actin machinery is inhibited by a mild Jasplakinolide treatment (Boulant *et al*., 2012; Kural *et al*., 2015) (Figure 2D, E).

**Figure 2.**
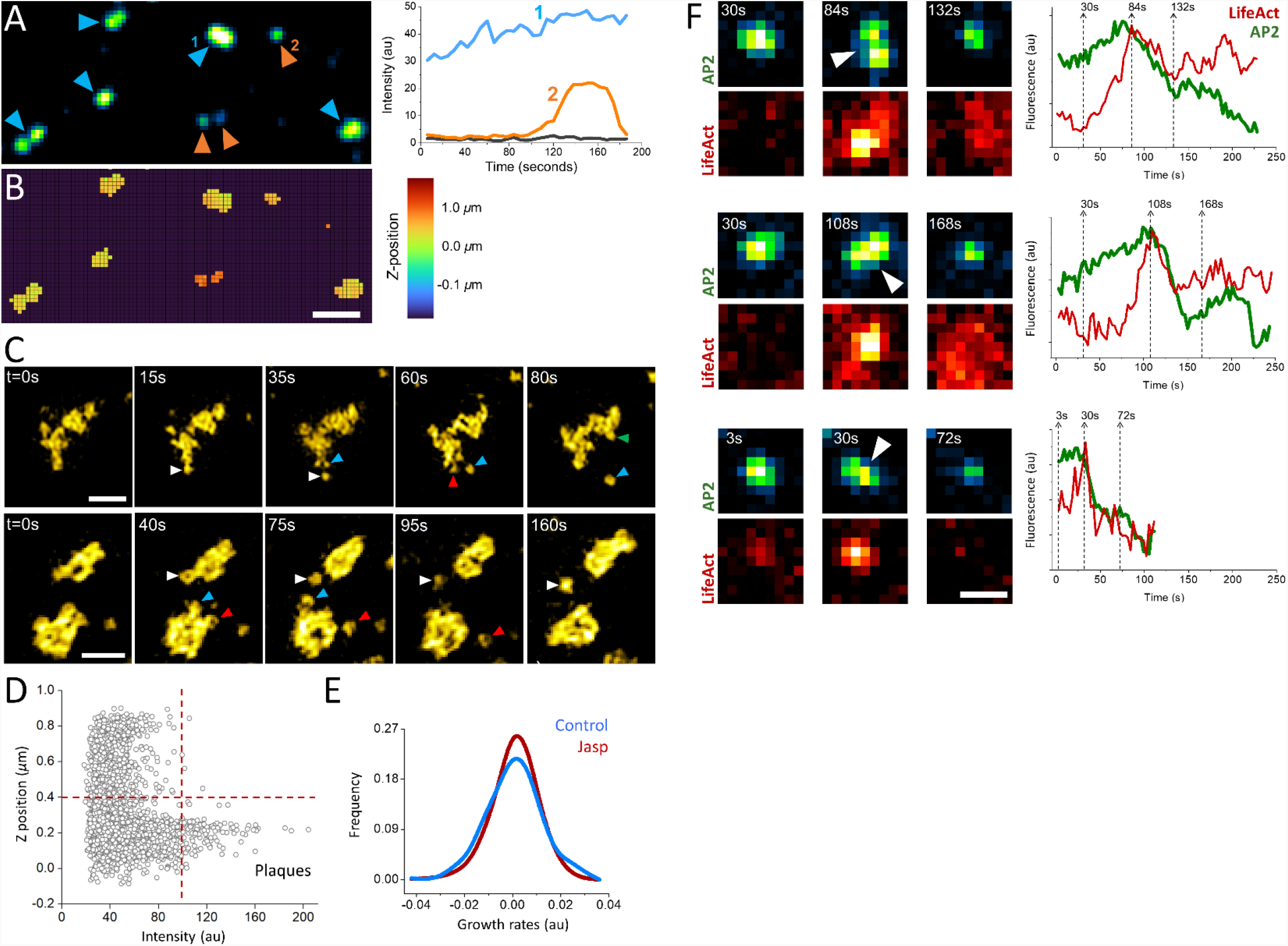
Role of actin in clathrin plaque dynamics. **(A)** Clathrin-coated pits and plaques are imaged on the ventral surface of live SUM-159 cells using spinning-disk confocal microscopy. The blue and orange arrowheads mark plaques and pits, respectively. Representative fluorescence intensities of a clathrin plaque (1) and pit (2) are shown on the left panel, where the black line shows the background intensity. Although its intensity has changed over time, the clathrin plaque was observable throughout the entire acquisition. Whereas the clathrin pit in the example had a lifetime of ∼100 seconds, it appeared around the 90^th^ second and was internalized at the end of the acquisition. **(B)** The average z-positions of the pits and plaques in (A) are determined for each pixel as described earlier (Kural *et al*., 2012). Plaques are localized to membrane-substrate contact sites and, therefore, are found in the same axial position. Whereas the clathrin pits are often observed at higher z positions (Ferguson *et al*., 2016). Scale bar, 1*µ*m. **(C)** Examples show endocytic pits (marked by arrowheads with distinct colors) budding off from large clathrin plaques. Scale bars, 1*µ*m. **(D)** Scatter plot shows the z-position versus the maximum intensity of clathrin-coated structures detected at the ventral surface of SUM-159 cells genome edited to express AP2-EGFP. The dashed lines denote the thresholds use to select plaques, which are brighter than pits and positioned closer to the substrate (Ferguson *et al*., 2016). **(E)** Growth rate distributions of plaques imaged in SUM-159 cell prior to (Control) and 10 minutes after treatment with 1*µ*M Jasplakinolide (Jasp). The narrower growth distribution upon Jasp treatment is a result of slower plaque dynamics upon inhibition of actin machinery (Ferguson *et al*., 2016). **(F)** Increased membrane tension upon cell squeezing gives rise to formation of large and bright clathrin puncta, which can split in multiple fragments that are internalized independently. Three such events are shown in both AP2 (to mark clathrin coats) and Lifeact (actin marker) channels, where the corresponding integrated fluorescence intensities are shown on the right. The dashed lines show the corresponding time points. The arrowheads mark the splitting events that coincide with a burst in actin recruitment that is followed by a significant reduction in the AP2 signal. Scale bars, 0.5 *µ*m.

Increased membrane tension is another factor that renders internalization of clathrin coats dependent on the forces generated by actin polymerization (Boulant *et al*., 2012; Kaur *et al*., 2014; Kaplan *et al*., 2022). When the plasma membrane tension is increased by squeezing cells using a soft polymer cushion, dynamic CCPs are replaced by bright fluorescent puncta corresponding to large and long-lived clathrin-coated structures as imaged under diffraction-limited fluorescence microscopy (Ferguson *et al*., 2017). We found that splitting events can be observed in this population of clathrin-coated structures and characterized by increased eccentricity of the fluorescent spot followed by internalization of a portion of the coat and a significant reduction in integrated fluorescence signal (Figure 2F). Splitting events coincided with a burst of actin fluorescence, suggesting that actin polymerization provides the force necessary for splitting of large clathrin coats into smaller pieces prior to internalization.

### Giant coated pits (GCPs) observed in tissues of developing fruit fly embryos

Despite its short penetration into the sample, TIRF-SIM can be used to monitor formation of individual clathrin-coated structures within tissues of developing *Drosophila* embryos expressing fluorescently tagged clathrin coat components (Willy, Ferguson, *et al*., 2021) (Video 1). In these *in vivo* acquisitions, we have detected formation clathrin plaques on migratory hemocytes that occasionally move adjacent to the vitelline membrane of the embryo (Figure 3A; Video 2). Similar to our observations in cultured cells (Figure 2C), we observed clathrin plaques detected *in vivo* also go through splitting events that give rise to smaller coat fragments internalized independently from the edge regions (Figure 3B).

**Figure 3.**
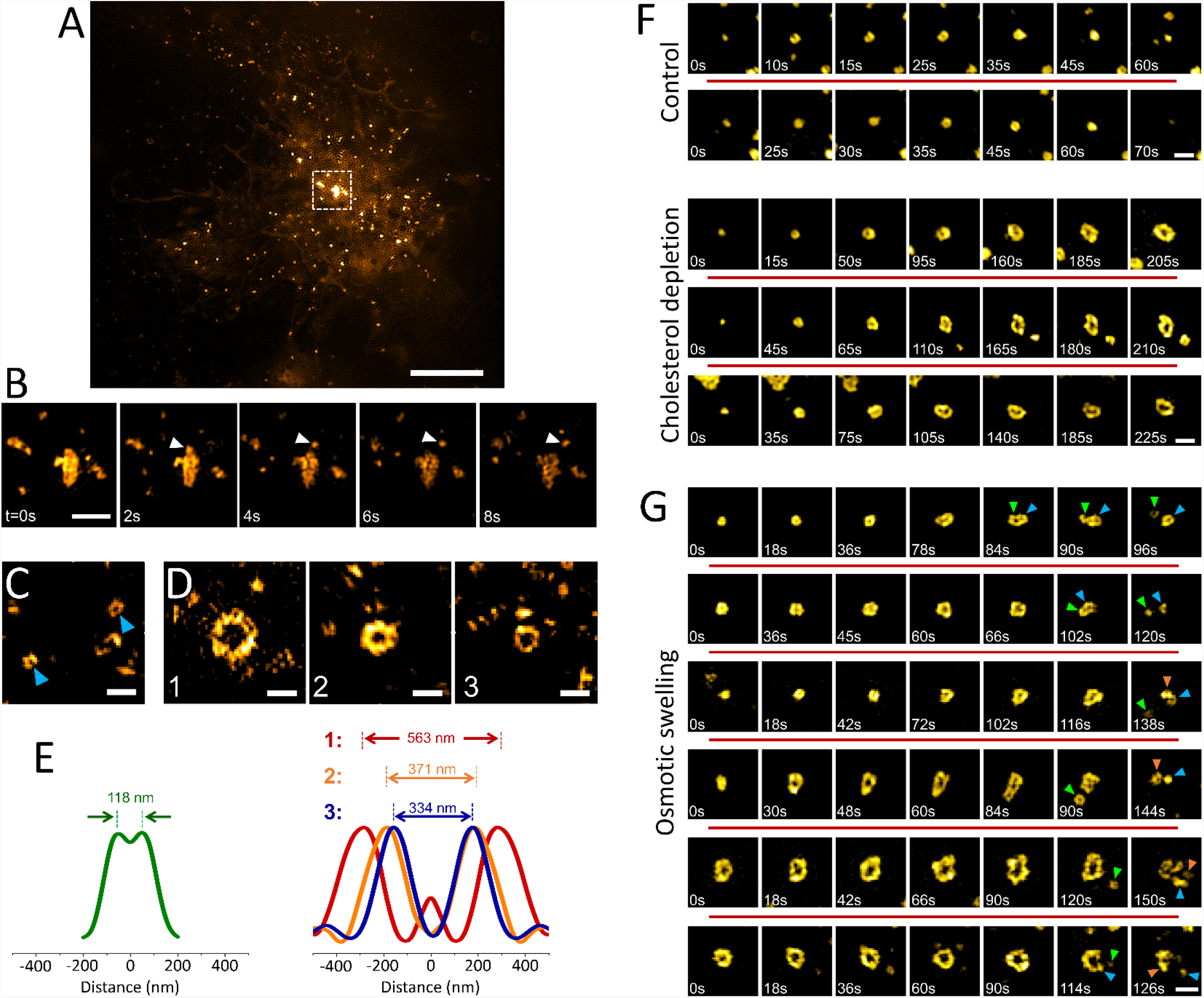
Formation and dissolution of giant coated pits (GCPs) living cells and tissues. **(A)** TIRF-SIM image of a *Drosophila* hemocyte within a live embryo. Scale bar, 5 *µ*m. **(B)** Montage shows the boxed region in (A). Arrowheads mark a coat fragment splitting from a clathrin plaque. **(C)** Arrowheads mark two canonical CCPs imaged within developing *Drosophila* embryos expressing clathrin-mEmerald. (**D)** GCPs imaged at the lateral epidermis tissue during dorsal closure of the embryo. **(E)** *Left:* Radial average of canonical clathrin pit images obtained from embryos, where the peak-to-peak distance is 118 nm. *Right:* Radial averages of the three GCPs shown in (D). The corresponding peak-to-peak separations are 563, 371 and 334 nm, respectively. **(F)** Examples (in different rows) of canonical CCP formation detected in live SUM159 cells expressing clathrin-mRuby (upper). Cholesterol depletion by methyl-β-cyclodextrin (MβCD) treatment induces GCP formation in the same cell line (lower). **(G)** Examples (in different rows) of GCP formation and dissolution detected upon hypotonic swelling in SUM159 cells genome edited to express AP2-EGFP. Arrowheads in different colors mark segments separated from the GCPs. Scale bars, 0.5 *µ*m.

Surprisingly, on the apical surface of epidermal cells of late *Drosophila* embryos, we have detected clathrin pits that are significantly larger than the canonical CCPs (Figure 3C, D; Video 2). The diameter of these structures reaches up to 0.5*µ*m, about 5-times wider than the typical size of CCPs observed within the same tissue (Figure 3E). We named these structures giant coated pits (GCPs) as their estimated volumes can be orders of magnitude larger than the canonical pits. Interestingly, a subset of GCPs were split into multiple fragments before internalization (Video 3). Considering the effects of high membrane tension on the size and curvature of clathrin-coated structures (Saias *et al*., 2015; Ferguson *et al*., 2017; Willy, Ferguson, *et al*., 2021), we hypothesized that the formation of GCPs is a result of tension building up on the lateral epidermis during late stages of embryogenesis (Kiehart *et al*., 2017).

### Increased membrane tension induces formation of GCPs in cultured cells

Membrane tension affects the dynamics and structure of endocytic clathrin coats by increasing the energy cost of curvature generation on the plasma membrane (Heuser, 1989; Subtil *et al*., 1999; Boulant *et al*., 2012; Ferguson *et al*., 2016, 2017; Willy *et al*., 2017; Willy, Colombo, *et al*., 2021; Willy, Ferguson, *et al*., 2021). Here, we employed two independent strategies to increase the membrane tension in cultured cells. First, we treated cells with methyl-β-cyclodextrin (MβCD) to deplete cholesterol from the plasma membrane (Ferguson *et al*., 2016; Willy *et al*., 2017; Biswas *et al*., 2019). Second, we used hypotonic medium to induce osmotic swelling in cells (Dai *et al*., 1998; Ferguson *et al*., 2017). We found that increased membrane tension due to cholesterol depletion gives rise to formation of GCPs in good agreement with our hypothesis (Figure 3F). Even though GCPs are substantially larger than the canonical CCPs and their formation takes markedly longer, the mechanism of curvature generation was the same: the footprint of the coat increased with the detection of the ring pattern, which indicates that the GCP curvature is generated at early stages of the coat formation without a flat-to-curved transition that requires restructuring of the coat (Willy, Ferguson, *et al*., 2021) (Figure 3F).

To induce osmotic swelling we exposed cells to 80% hypotonic medium (Willy, Colombo, *et al*., 2021). In contrast to cholesterol depletion, osmotic swelling and its effects on the plasma membrane tension and clathrin-mediated endocytosis dynamics is temporary (Ferguson *et al*., 2017). We observed GCP formation within the first 15 minutes of the hypo-osmotic treatment, where the membrane tension is expected to be the highest (Bucher *et al*., 2018; Willy, Ferguson, *et al*., 2021). As in *Drosophila* embryos, we found that GCPs observed in swollen cells split into multiple small pieces, some of which have the characteristic ring pattern of the canonical CCPs (Figure 3G).

## DISCUSSION

In this study, we developed and used advanced super-resolution imaging modalities to study the formation and internalization dynamics of distinct classes of endocytic clathrin-coated structures within cultured cells and tissues of developing *Drosophila* embryos. We have focused on three different mechanisms of clathrin-coated vesicle formation: in addition to canonical CCPs that form *de novo*, and CCPs internalized from the edges of plaques, for the first time, we report a new mechanism where endocytic vesicles form by dissociation from giant coated pits (GCPs) (Figure 4A-C). GCPs can be orders of magnitude larger than the canonical pits and have longer lifetimes, however, their mechanism of curvature generation is the same.

**Figure 4.**
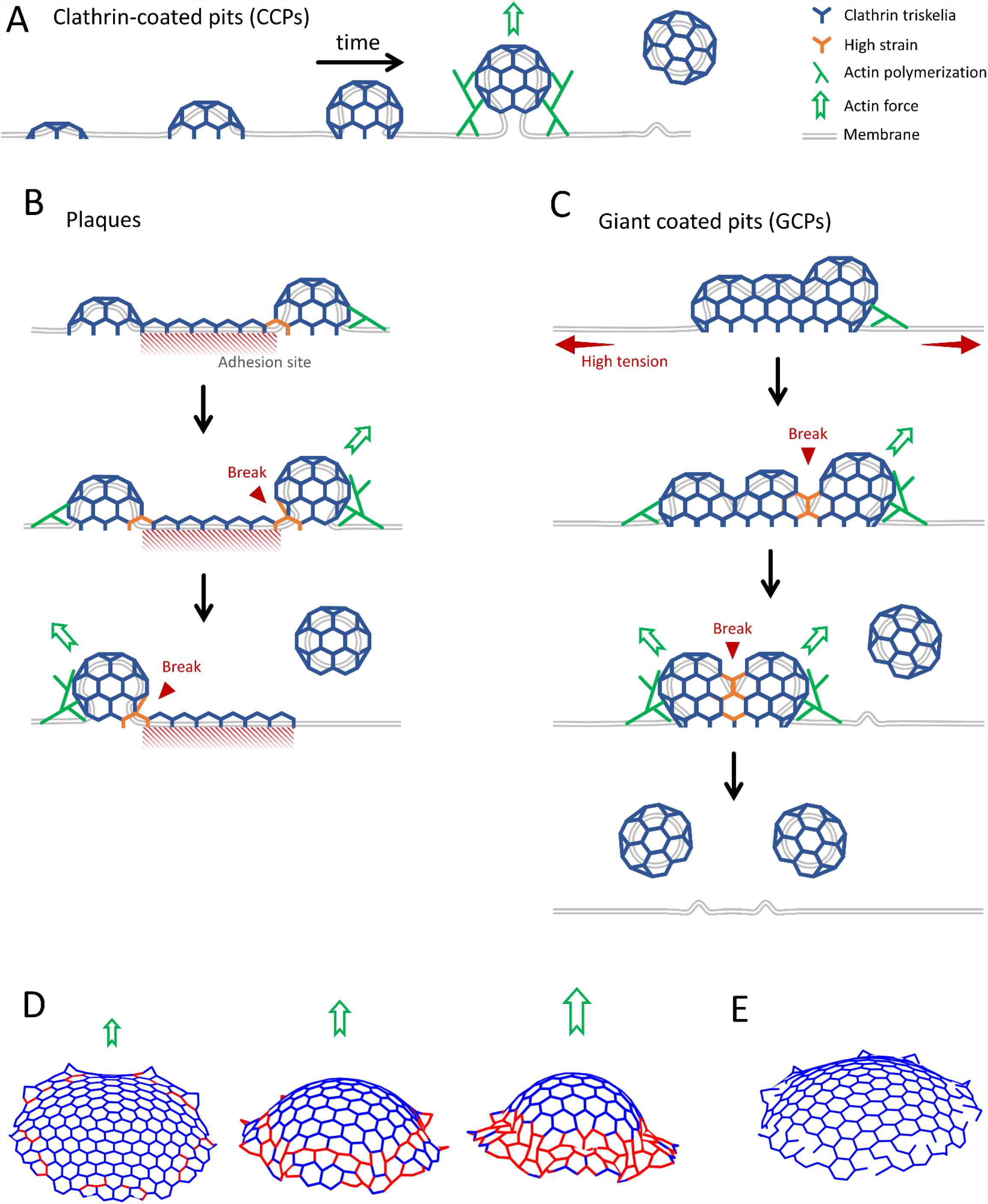
Distinct mechanisms of internalization of endocytic clathrin-coated vesicles. **(A)** Cartoon representation of growth and internalization of canonical CCPs. The inward movement of the pit during internalization is dependent on actin dynamics, which becomes indispensable under increased membrane tension (Boulant *et al*., 2012; Ferguson *et al*., 2017). **(B)** Clathrin plaques are predominantly observed at the membrane-substrate adhesion sites. It was proposed that clathrin pits can break apart from the edges of plaques and grow into coated vesicles independently. This “rupture and growth” model proposes that increased local curvature creates strain on the lattice, giving rise to breaks and splitting coat fragments (den Otter and Briels, 2011; Lampe, Vassilopoulos and Merrifield, 2016; Willy, Ferguson, *et al*., 2021). **(C)** GCPs form under increased membrane tension. We propose that forces produced by actin polymerization, aided by reducing plasma membrane tension, give rise to splitting of GCPs into multiple small fragments that are internalized independently. **(D)** Snapshots of simulations showing how a low curvature coat transforms into a dome with increasing force. The elastic energy of red triskelions is much higher than the blue ones. The lattice can break at the location of red triskelions if the strain is too high **(E)**.

We found that, unlike osmotic swelling experiments, GCPs forming upon MβCD treatment were stalled and did not undergo splitting. Note that cholesterol depletion and osmotic swelling increase the effective membrane tension in different ways. Cholesterol depletion increases the adhesion of actin cytoskeleton to the plasma membrane (Khatibzadeh, Nima, Gupta, 2012), which may negatively impact actin dynamics. Whereas osmotic swelling increases the in-plane membrane tension without affecting the function of actin machinery (Dai *et al*., 1998; Boulant *et al*., 2012; Ferguson *et al*., 2017; Kaplan *et al*., 2022). Considering the fact that actin machinery becomes indispensable for internalization of clathrin-coated vesicles under high mechanical load (Boulant *et al*., 2011; Ferguson *et al*., 2017), we propose that splitting and internalization of GCPs under high membrane tension is also dependent on the forces provided by actin polymerization. Similar to the ‘rupture and growth” model proposed for plaques (Figure 4B), local changes in the GCP curvature imposed by actin polymerization is expected to increase the strain and give rise to breaks on the coat (Figure 4C) (den Otter and Briels, 2011; Lampe, Vassilopoulos and Merrifield, 2016; Willy, Ferguson, *et al*., 2021). To test how increasing curvature would affect strain on the coat, we simulated the transformation of a clathrin lattice containing a pentagonal defect to a dome using a steepest descent technique, in which we calculated the total energy of each vertex and moved the vertices as follows: 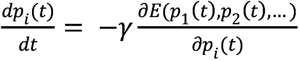 where *p* is the position of each vertex, *γ* the relaxation rate, and *E* the energy of each vertex. The total energy depends on the state of triskelions: stretched, compressed and angle between them (see the materials and methods section for the details). Figure 4D shows snapshots of how the lattice distorts as the strength of the force that induces curvature on the complex increases. If the energy of an edge exceeds a threshold value (see the materials and methods), the edge is colored red. Otherwise, it is colored blue. As suggested by the “rupture and growth” model, the deformations resulting at the edges of the coat lead increased strains in these regions, which may result in breaks in the lattice leading to fragmentation of the complex (Figure 4E) and, subsequently, growth and internalization of CCPs independently.

As mentioned before, GCP formation is observed within the first 15 minutes of the hypo-osmotic treatment. After this point, membrane tension is expected to converge back to original levels as osmotic swelling is temporary and endocytic clathrin dynamics recover from the inhibitory effects of increased membrane tension within 30 minutes after the hypotonic treatment (Ferguson et al., 2017b; Willy, Ferguson, et al., 2021). Forces provided by actin filaments may increase the strain on the coat more efficiently if the underlying membrane ceases to support the rigidity of the coat. On the other hand, such a recovery does not take place after cholesterol depletion, which may be another reason why GCP splitting events were not observed in MβCD treated cells.

Finally, our *in vivo* data acquired from developing *Drosophila* embryos suggest that GCPs are physiologically significant. On the other hand, in comparison to the canonical CCPs, formation of GCPs is very sparse (∼4.5 ×10^−3^/*µ*m^2^), suggesting that they only form under extreme mechanical conditions such as increased membrane tension. Considering the effects of the cargo size on the geometry and dynamics of endocytic clathrin-coated structures, we expect clustering of ligands, or their receptors may also give rise to formation of GCPs.

## MATERIALS and METHODS

### Fluorescence live cell imaging

Genome-edited AP2-eGFP expressing human breast cancer SUM159 cell lines were a kind gift of Dr. Tomas Kirchhausen (Harvard Medical School). Cells were grown at 37 °C and 5% CO_2_. SUM159 cells complete media consisted of F-12/Glutamax (Thermo Fisher Scientific), supplemented with 5% fetal bovine serum (Gibco), 100 U/mL penicillin and streptomycin (Thermo Fisher Scientific), 1 μg/mL hydrocortisone (H-4001; Sigma-Aldrich), 5 μg/mL insulin (Cell Applications), and 10 mM 4-(2-hydroxyethyl)-1-piperazine-ethane-sulfonic acid (HEPES), pH 7.4. Complete L15 media (Thermo Fisher Scientific) supplemented with 5% serum and 100 U/ml penicillin and streptomycin was used during live-cell imaging, as it supports cell growth in environments without CO_2_ equilibration. In actin inhibition assays, cells were treated with 1 μM Jasplakinolide (Alexis Biochemical) for 10 minutes and imaged with a spinning disk confocal microscopy setup built on a Eclipse TI-E microscope (Nikon Instruments Inc.) equipped with a perfect focusing system (PFS), a temperature-controlled chamber, CSU-W1 spinning disc unit (Yokogawa Electric Corporation), a 100× objective lens (Nikon CFI Plan-Apochromat Lambda, NA 1.45), an Electron Multiplying Charge Coupled Device (EMCCD) camera (iXon DU897 Ultra; Andor Technology), and 488-and 640-nm excitation lasers with 100 mW of nominal power. Images were acquired at a rate of 0.25–0.5 Hz with a laser exposure of 100 ms per frame. Image acquisition was done using NIS Elements software.

### Structured Illumination Microscopy (SIM)

The schematic of the home-built high NA SIM system is shown in Supplementary Figure 1. The beams from 488 nm and 561 nm (300mW, Coherent, SAPPHIRE LP) lasers are collinearly combined. An acousto-optic tunable filter (AOTF; AA Quanta Tech, AOTFnC-400.650-TN) is used to switch between them and to adjust the illumination power. After passing through the AOTF, the beam is expanded and sent to a phase-only modulator (Kner *et al*., 2009), which is used to diffract the beam. The phase-only modulator consists of a polarizing beam splitter, an achromatic half-wave plate (HWP; Bolder Vision Optik, BVO AHWP3), and a ferroelectric spatial light modulator (SLM; Forth Dimension Displays, QXGA-3DM-STR).

For high NA SIM acquisitions, nine grating patterns consisting of 3-orientations and 3-phases are displayed on the SLM. The diffracted beams then pass through an azimuthally patterned achromatic half-wave plate (Azimuthal HWP; Bolder Vision Optik), which consisted of three pairs of segments with custom designed fast axis orientations and the linear polarization of the diffracted light from the grating patterns is rotated to the desired s-polarization (Guo *et al*., 2018). The beams then pass through a mask in order to filter out undesired diffractions and pass only ± 1 diffraction orders, which are then relayed onto the back focal plane of the high numerical aperture (NA) objective (Olympus, APO 100X 1.65 OIL HR 0.15) mounted on an inverted Eclipse TI-E microscope (Nikon instruments Inc.). The incidence angle onto the interface of the specimen and coverslip (V-A Optical Labs, SF-11) is adjusted by changing the period of the pattern displayed on the SLM.

For each frame two SIM acquisitions are taken at two incidence angles which are slightly lower (grazing incidence illumination) or higher (TIR illumination) than the critical angle. The fluorescent emission generated by the applied excitation pattern of each phase and orientation is collected by the same objective and focused by a tube lens onto an EMCCD camera (Andor, iXon Ultra 897). The acquired nine raw images are reconstructed into a super-resolution image based using a previous algorithm (Wiener coefficient = 0.001) (Huang *et al*., 2018). The cells were left in the incubator for 3 hours to allow complete spreading before starting imaging using 488 nm illumination at 2 frames/sec with 20 msec acquisition time.

TIRF-SIM imaging assays of live *Drosophila* embryos and SUM159 cells treated with hypotonic shock and cholesterol depletion were conducted at the Advanced Imaging Center (AIC) of the Janelia Research Campus using a lower NA objective (Olympus UAPON 100X 1.49 OTIRF) as described before (Willy, Ferguson, *et al*., 2021). This system was built using an inverted microscope (Axio Observer, ZEISS) equipped with a sCMOS camera (Hamamatsu, Orca Flash 4.0). Images were acquired at rates ranging from 2s/frame to 6s/frame using 20ms exposure time.

### 3D tracking of clathrin-coated pits and plaques

For each frame in the 3D time-lapse movie, max intensity value is picked out of all z stacks at each x-y position to form a 2D movie that contains the maximum intensity. The nuclei traces are then tracked using the TraCKer algorithm (Ferguson *et al*., 2016) on the 2D maximum intensity movie. TraCKer uses an algorithm based on Mexican hat function to detect traces. For each time point in detected traces, axial (z) position is determined by calculating the center of mass across all z stacks at tracked x-y position. A trace-wide axial position is assigned by calculating the mean z position of all time points in the trace.

Growth rates (slopes) are extracted from the intensity values on the tracked traces as described before (Ferguson *et al*., 2016). Trace intensities are normalized by subtracting a constant background value. Traces with lower than 12s are discarded. For each 12s interval, a least-square fit is applied to determine the slope at each time point of the trace.

### Simulations of clathrin-coated structures

We start with a clathrin complex consisting of a pentagon surrounded by a honeycomb lattice, as shown in Supplementary Figure 2.

The lattice as shown lies in a plane. We then introduce a set of terms contributing to overall energy that, when minimized, leads to the deformation of the lattice into a dome-like structure. The contributions to the energy of the complex are

1. The energy associated with the angle between adjacent edges of a hexagon
2. The energy associated with the length of a bond
3. The “pucker angle” energy
4. The tethering energy

We address each energy separately below

#### 1. The energy associated with adjacent edges of a hexagon

Consider a given vertex and the three edges that attach to it, as shown in Supplementary Figure 3.

If the edges lie in a plane, and all the angles are equal, then 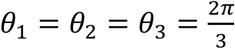. However, if all angles are the same and are less than 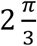, then the vertices cannot lie in a plane. In this case the resulting configuration looks like one of the vertices and three of the edges of a tetrahedron. The energy that favors a departure from planar geometry is introduced via the scalar product of adjacent edges attached to a vertex. Let *ê*_*i*_ (1≤*i*≤ 3) be the three unit vectors parallel to the edges incident on a given vertex with tails on the vertex. We construct the energy so that it favors a configuration in which *ê*_*i*_· *ê*_*j*_ = cos(*θ*_0_), with *θ*_0_ < 2 *π*/3. We adopt the following form for this energy,

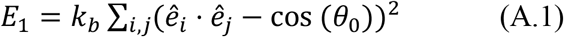

where the sum is over the nearest neighbor edges. The larger the bending modulus *k*_*b*_ is the more important the “angular’” energy.

#### 2. The energy associated with the lengths of the edges

In the case of the honeycomb lattice in a plane all edges have the same length. When the lattice acquires a curvature, it becomes necessary to add an energy consistent with the desired lengths of the edges. We will assume a preferred length of *l*_0_. We introduce an energy favoring that length through the expression

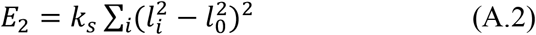

where the sum is over link index *i*, and *l*_*i*_is the length of the link number *i*. Again, the larger the stretching modulus *k*_*s*_, the more important this contribution to the total energy to be minimized.

#### 3. The “pucker angle” energy

This energy is based on the volume associated with a local curvature. Consider the triplet of edges shown in Supplementary Figure 3. We can think of this as a projection of three sides of a parallelepiped, shown in Supplementary Figure 4.

If the three edges incident on the vertex shown as a dot in the figure are 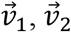 and 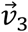, then the volume of the parallelepiped is equal to 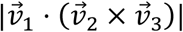. An important feature of the triple product is that with the absolute value lines removed, it can be positive or negative. Specifying a value of the triple product gives us a set of four vertices that produce a positive or negative curvature. If the three edges are coplanar, then the triple product, and hence the curvature, are zero. It is possible then to introduce an energy based on the triple product favoring a specific local curvature.

Calling this triple-product-based curvature 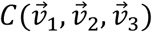, or *C*_*i*_for short, the energy is of the form

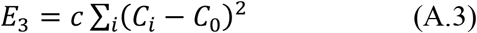

where *C*_0_ is the preferred value of the triple product. This is directly related to the pucker angle in (den Otter and Briels, 2011). The positive coefficient *c* controls the importance of this contribution.

#### 4. The tethering energy

The final energy is related to the attraction between the clathrin complex and the plasma membrane. Since the membrane is under tension, the clathrin lattice embedded in the membrane cannot bend and form a shell. We consider the forces keeping the clathrin lattice flat to be Hooke’s law like. If the coordinate of each vertex is 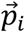 and the membrane is in the *x*-*y* plane, the total tethering energy is proportional to

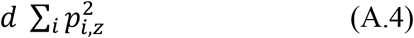

The importance of this energy follows from the magnitude of the positive coefficient *d*.

As the tension in the membrane decreases, the magnitude of *d* decreases. The actin network could also change the amplitude of this force.

##### The minimization procedure

We use a straightforward steepest descent method. Given a parameter-dependent energy of the form *E*(*p*_1_, *p*_2_, …) we introduce a time-dependence, so that *p*_*i*_→ *p*_*i*_(*t*). Then, we solve the equations of motion

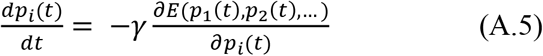

##### Evolution of the clathrin complex and bond breaking

The overall idea is that when hexagons—and the single pentagon—become sufficiently distorted, then bonds can be broken. The mechanism for determining when this happens depends on the angles between adjacent triskelion bonds. See Supplementary Figure 5.

Note that there are up to four relevant angles between adjacent vertices to consider in determining whether or not the central bond breaks or remains intact. The basis for the determination of whether or not a bond is broken is the expression on the right hand side of (A.1), with the coefficient *k*_*b*_ set equal to one and the preferred angle *θ*_0_ set equal to 2*π*/3, so cos(*θ*_0_) = −0.5. For each central bond, shown in red we calculate the sum of the two, three or four expressions, depending on the number of external bonds, shown in black. If that sum exceeds a chosen “breakpoint” value, the bond is deemed to have broken. Otherwise, the bond remains intact.

As an example of the application of this analysis, we have run a simulation of the development of a dome-like structure in a clathrin complex originally confined to a plane, as shown in Figure 4D in the paper.

At any point in the process, as shown in Figure 4E we can assess the bonds for removal through breakage. Note that the intact hexagons, and the central pentagon, are relatively undistorted.

## ACKNOWLEDGEMENTS

We thank John Heuser for helpful discussion at the early phase of this project. We also thank the Advanced Imaging Center (AIC) at Janelia Research Campus for access to their TIRF-SIM system. We are particularly indebted to Aaron Taylor and Satya Khuon from the AIC team. The AIC is jointly supported by the Howard Hughes Medical Institute and the Gordon and Betty Moore Foundation. C.K. was supported by NIH R01GM127526 and NSF Faculty Early Career Development Program (award number: 1751113). E.C. was partially supported by the Pelotonia Young Investigator Award, IRP46050-502339. R.Z. is supported by NSF grant DMR-2131963.

## AUTHOR CONTRIBUTIONS

C.K. conceived the study. A.A. and C.T. developed the home-built TIRF-SIM setup and performed the SIM measurements at the TIRF and GI modes. C.K. and E.C. performed the TIRF-SIM experiments on *Drosophila* embryos and SUM-159 cells involving hypotonic shock and cholesterol depletion at AIC. U.D. has designed and performed experiments involving actin inhibition. T.W. performed the three-dimensional tracking and growth rate distribution analyses. R.Z. and J.R. simulated the strain on clathrin coats upon curvature increase. C.K. wrote the manuscript.

## SUPPLEMENTARY MATERIALS

**Supplementary Figure 1.**
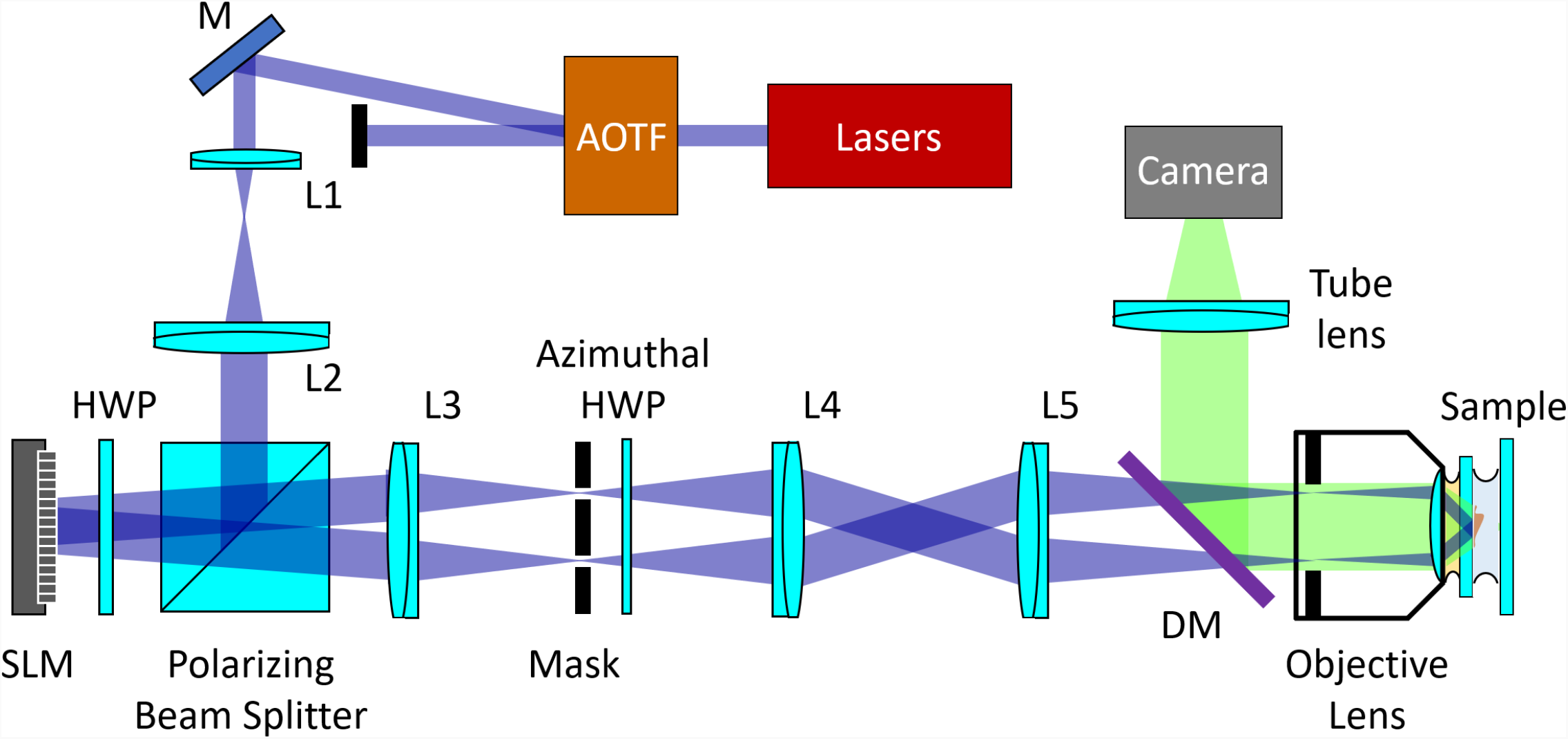
Optical configuration of the home-built structured illumination microscopy (SIM). AOTF: acousto-optic tunable filter; HWP: half wave plate; SLM: spatial light modulator; L1-L5: lenses; M: mirror; DM: dichroic mirror.

**Supplementary Figure 2.**
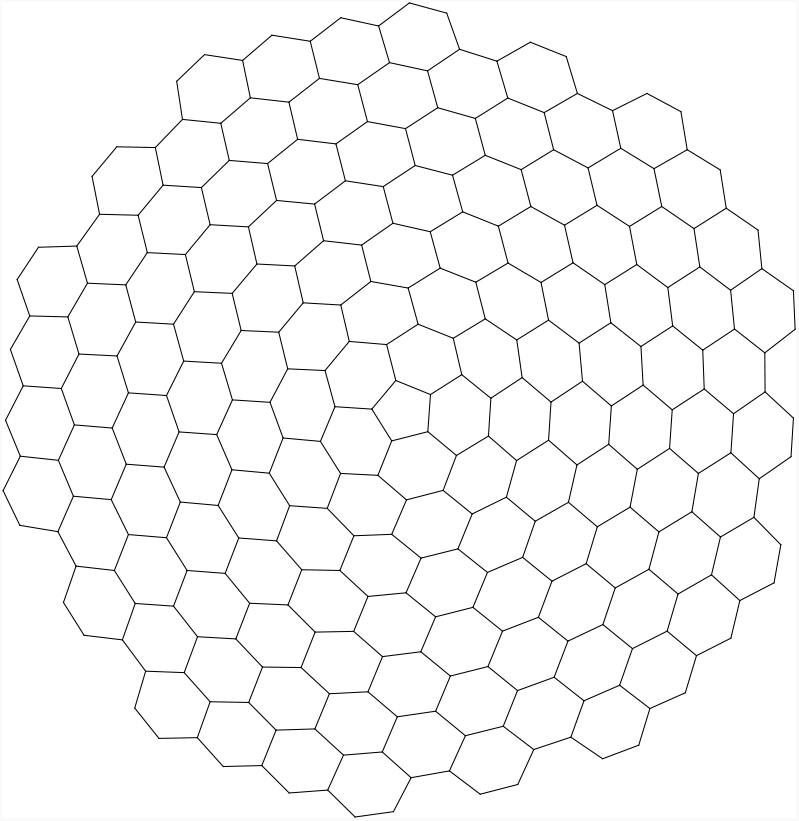
The initial complex, consisting of a pentagon surrounded by a honeycomb-like lattice. The complex lies in a plane, which means that the hexagons are distorted versions of the regular hexagons in a true honeycomb lattice.

**Supplementary Figure 3.**
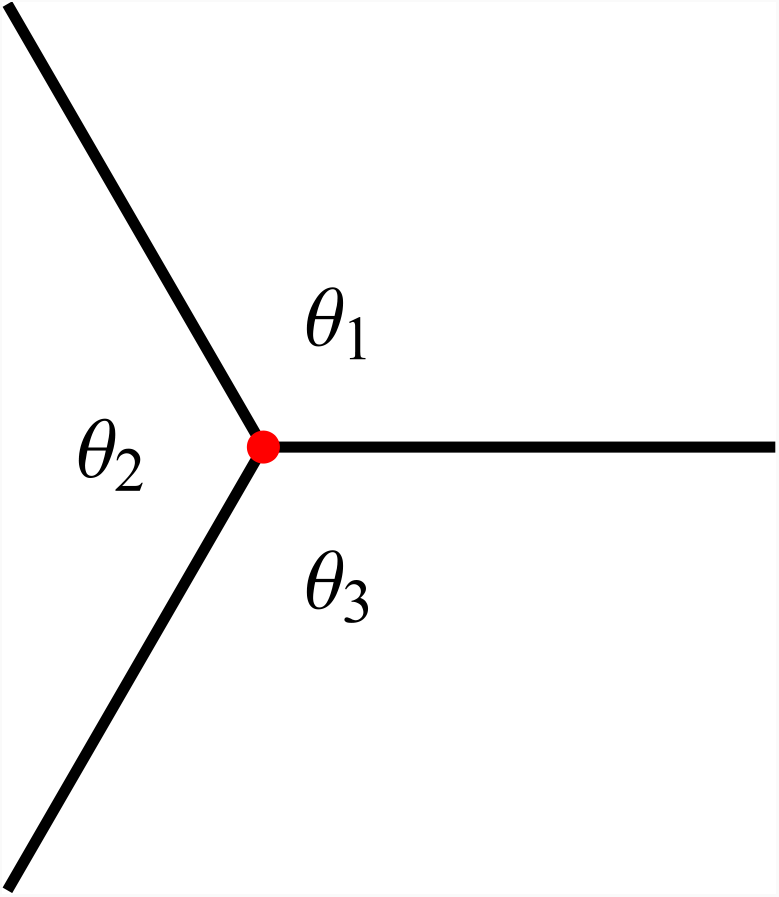
A vertex and the three edges attached to it, along with the angles between the edges.

**Supplementary Figure 4.**
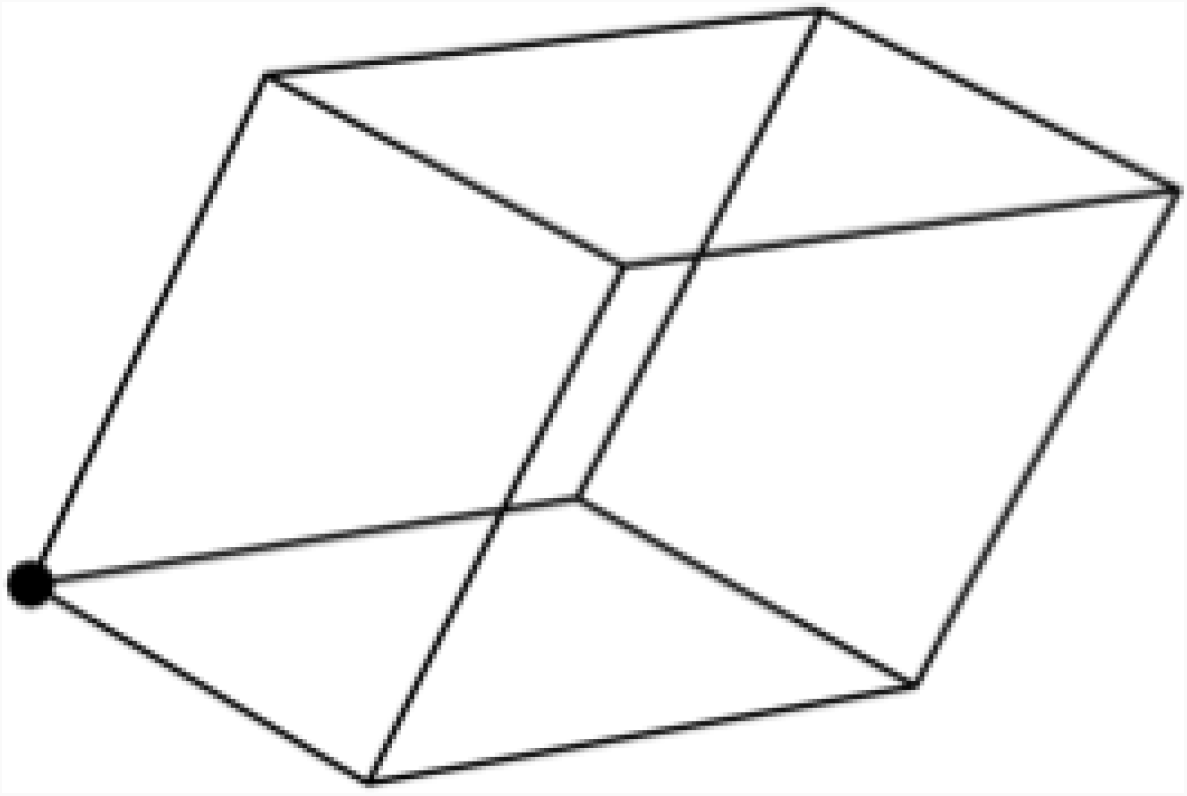
The parallelepiped constructed from the three edges incident on the vertex shown as a dot.

**Supplementary Figure 5.**
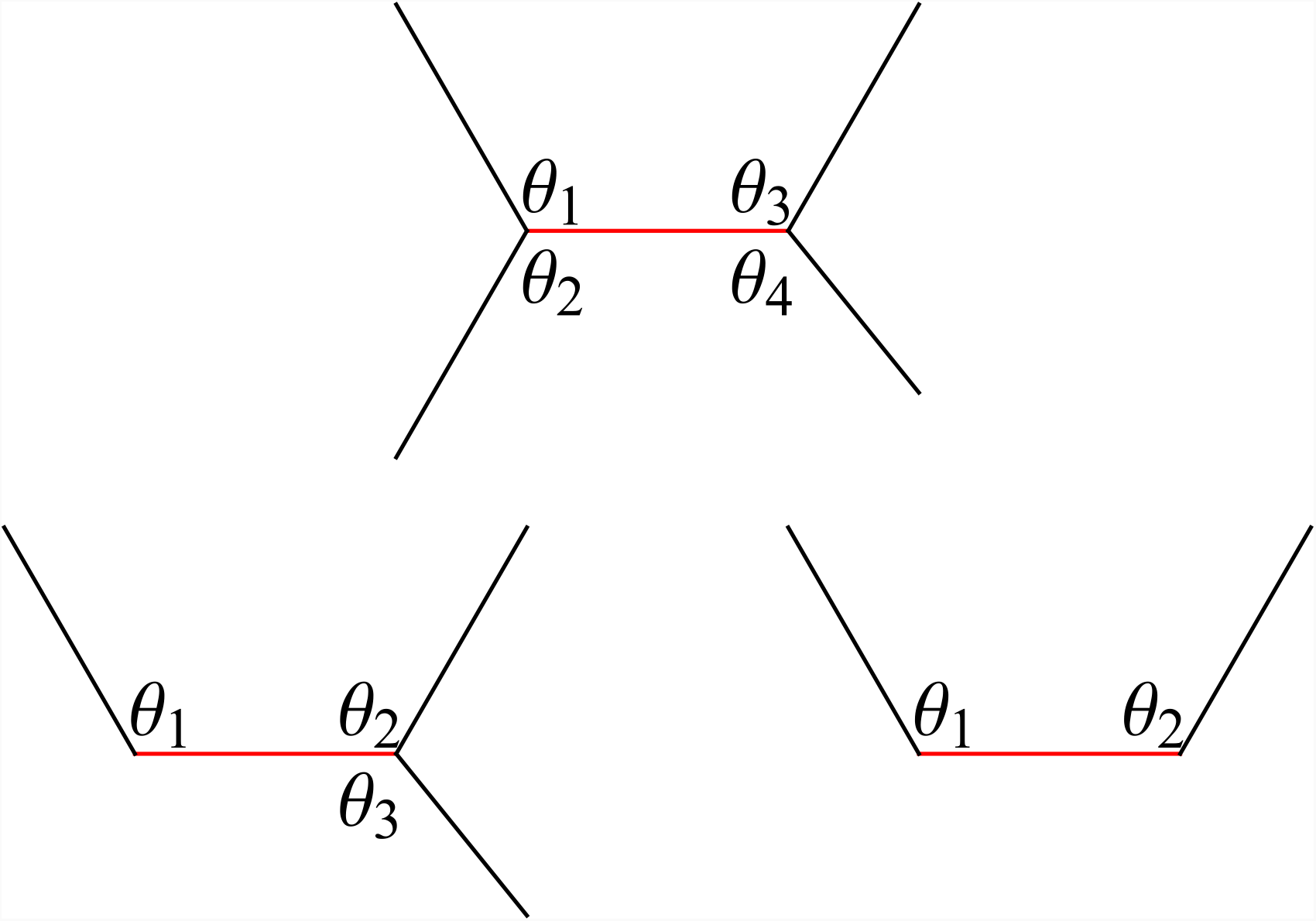
The central bond, in red, joins two nearest neighbor vertices. The two vertices have either one or two additional bonds attached. The angles between adjacent bonds are *θ*_1_, … *θ*_4_, as shown.

**Supplementary Video 1.** TIRF-SIM movie of clathrin coat dynamics acquired at the amnioserosa tissue of a late *Drosophila* embryo expressing clathrin-mEmerald. Further zoom-in (blue box) shows growth and dissolution of an individual clathrin-coated pit (arrowhead).

**Supplementary Video 2.** TIRF-SIM movie of clathrin coat dynamics acquired at the lateral epidermis of a late *Drosophila* embryo expressing clathrin-mEmerald. The panels below are the zoom-ins corresponding to four GCPs. The 4^th^ GCP gets out of focus due to a hemocyte entering the field of view and migrating between the epidermis and the vitelline membrane.

**Supplementary Video 3.** Examples of GCPs splitting into multiple fragments at the lateral epidermis of late Drosophila embryos.

## Notes

### Competing Interest Statement

The authors have declared no competing interest.

